# Mathematical model for rod outer segment dynamics during retinal detachment and reattachment

**DOI:** 10.1101/2024.01.08.574604

**Authors:** William Ebo Annan, Emmanuel O.A. Asamani, Diana White

## Abstract

Retinal detachment (RD) is the separation of the neural layer from the retinal pigmented epithelium thereby preventing the supply of nutrients to the cells within the neural layer of the retina. In vertebrates, primary photoreceptor cells consisting of rods and cones undergo daily renewal of their outer segment through the addition of disc-like structures and shedding of these discs at their distal end. When the retina detaches, the outer segment of these cells begins to degenerate and, if surgical procedures for reattachment are not done promptly, the cells can die and lead to blindness. The precise effect of RD on the renewal process is not well understood. Additionally, a time frame within which reattachment of the retina can restore proper photoreceptor cell function is not known. Focusing on rod cells, we propose a mathematical model to clarify the influence of retinal detachment on the renewal process. Our model simulation and analysis suggest that RD stops or significantly reduces the formation of new discs and that an alternative removal mechanism is needed to explain the observed degeneration during RD. Sensitivity analysis of our model parameters points to the disc removal rate as the key regulator of the critical time within which retinal reattachment can restore proper photoreceptor cell function.

## Introduction

The retina is the part of the eye containing photoreceptor cells [1]. In vertebrates, photoreceptor cells are classified into two, rods and cones [2]. Structurally, rod and cone cells have outer segment (OS), inner segment (IS), cell body, and synaptic terminals [3] (see Fig 1A). The OS of these cells is filled with tiny membranous discs harboring visual pigment molecules responsible for capturing and converting light energy into electrical signals perceived by the brain to enable vision [4–9]. Rod and cone cells differ in the shape of their OS, in functionality, and the nature of the discs comprising their OS. Morphologically, the cone OS has a cone-like shape with continuous discs connected to the OS cell membrane and are opened to the extracellular space. Cones are less sensitive to light and are responsible for color vision. On the other hand, rod cells have a cylindrical OS, with discrete discs that are separated from each other and the surrounding plasma membrane. They are highly sensitive to light and are responsible for night vision [3, 10]. The retina has two main layers, namely, the retinal pigmented epithelium (RPE) and the neural layer (NL). The NL of the retina contains the photoreceptor cells as well as bipolar, horizontal, amacrine, ganglion, and muller glial cells which work together to transmit signals to the brain to enable vision [11]. The RPE is a single-layer epithelium tightly joined to form a barrier between the retina and the choroid, a vascular structure surrounding the retina that supplies nutrients to the inner part of the eye and aids in thermoregulation [12]. The RPE contains lysosomes which break down discs that are shed from the rod and cone OS. It contains a dark pigment known as melanin which absorbs extra rays of light that can blur image formation and also prevents blood from entering the inner part of the eye [11].

**Fig 1.**
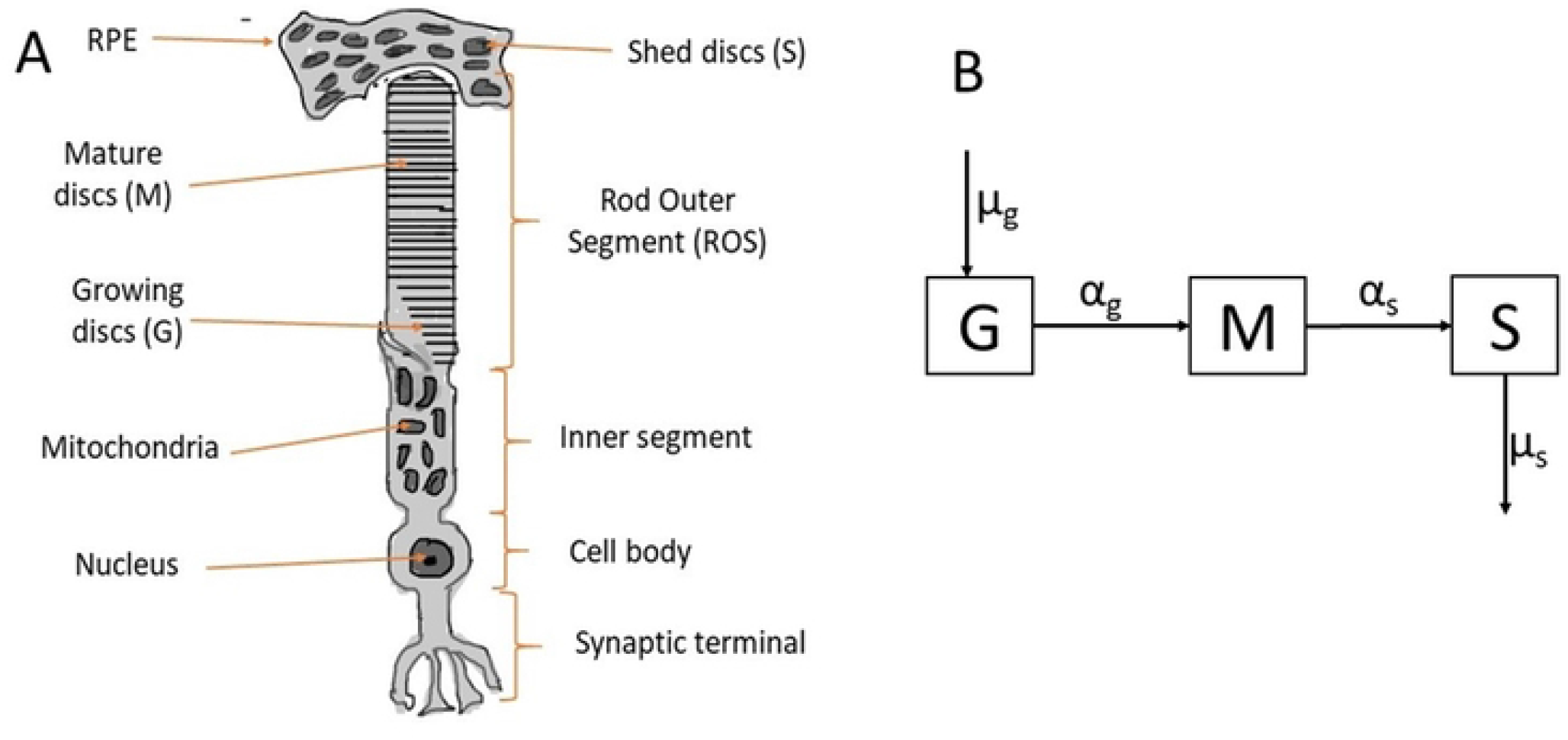
Rod cell and compartmental diagram. **A** shows the four main parts of a rod cell, namely the rod outer segment (ROS), inner segment, cell body and the synaptic terminal. In addition, we highlight the RPE where shed discs are disposed of. The categorization of discs in various compartments is also shown. **B** shows the transition of discs from one compartment (growth, mature and shed) to another. Discs are added to the G compartment at rate *μ*_*g*_, they mature to the M compartment at rate *α*_*g*_, are shed from the M compartment at rate *α*_*s*_, and are disposed of in the S compartment at rate *μ*_*s*_.

**Fig 2.**
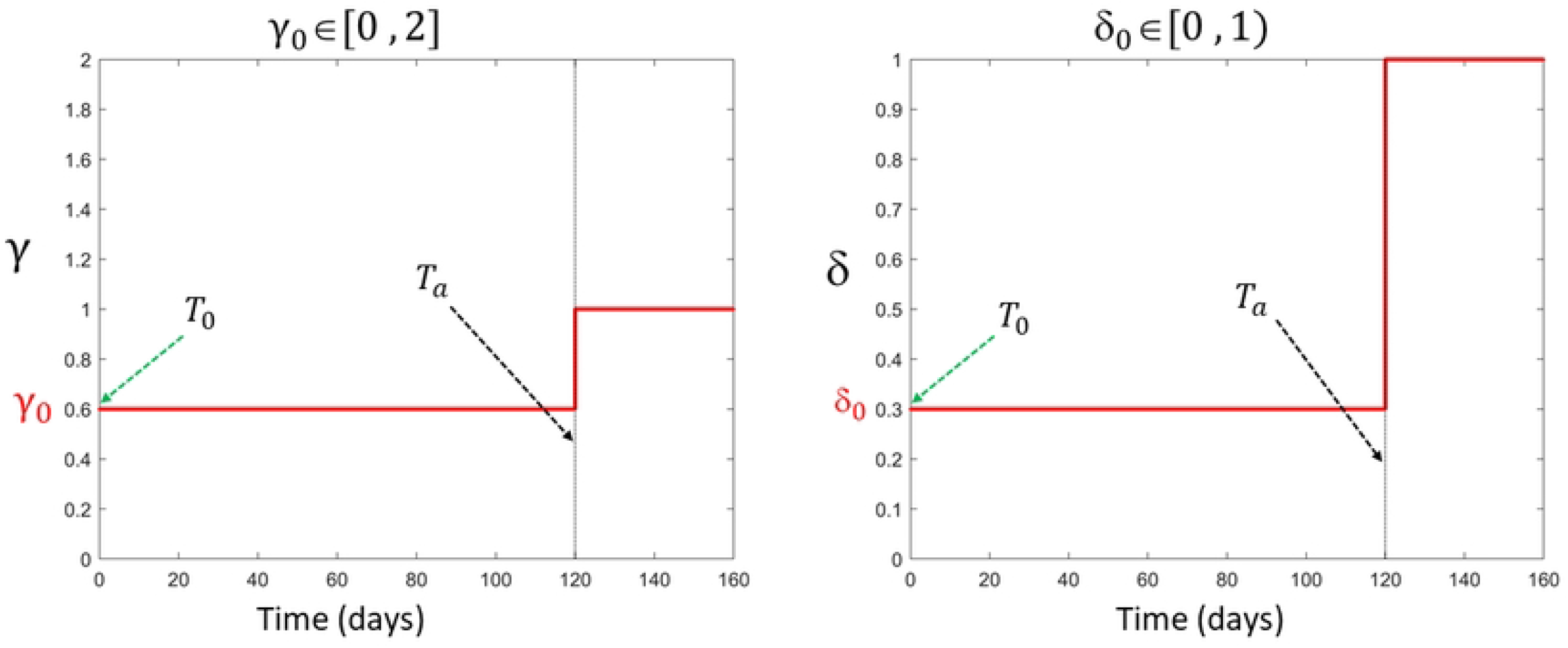
Step functions gamma and delta. Illustration for functions *γ*(*t*) (which affects disc removal rate) and *δ*(*t*) (which affects the discs addition rate), that take on values *γ*_0_ ∈ [0, 2] and *δ*_0_ ∈ [0, 1) respectively, during RD. Setting both parameters equal to 1.0 signifies normal cell behavior, and hence reattachment of the retina to the RPE at the time *T*_*a*_, (here 120 days).

Rods and cones undergo daily renewal of their OS through the addition of new discs at their base and the shedding of older discs from their distal ends to prevent accumulation of toxins caused by photo-oxidative compounds [13, 14]. Retinal detachment (RD), a separation of the NL from the RPE disrupts this normal renewal process. Factors such as old age, eye inflammatory diseases, tumors in the eye, or injury/trauma to the eye can cause the retina to detach [15–18]. When the retina detaches, the photoreceptor OS begins to degenerate/shorten [19, 20], yet the dynamics of this shortening are not well understood. For instance, some studies suggest that RD affects the renewal process by halting disc addition at the base of the OS [19] whereas other studies suggest that the addition of new disks continues during RD [16, 21]. It is however well established that shedding, as understood by the engulfment of discs by the RPE, stops during RD due to the separation of the NL from the RPE [19, 20]. The photoreceptor cells in a detached retina die after a certain period [16, 17, 21–23], however, if RD is treated on time, and the retina is reattached to the RPE, the possibility of rod and cone cell regeneration, and hence restoration of vision is high. Currently, the time frame within which retinal reattachment must be performed to restore vision is unknown [23].

To our knowledge, there are no mathematical models that explore the dynamics of the photoreceptor OS renewal during RD. Here, we extend the mathematical model we developed to study the normal dynamics of rod outer segment (ROS) renewal [24] to investigate the effect of retinal detachment on the renewal process. We focus our study on rod cells since the discrete nature of discs in rods, as opposed to the continuous nature of discs in cones, makes it easy to track the location of discs over time. Such tracking enables the estimation of the length of the ROS for validating mathematical models (as explained in Section Conversion from disk number to length). Another reason to study rod cells is that the survival of cone cells depends on a certain type of protein known as rod-derived cone viability factor (RdCVF) produced by rod cells [25]. It is typically the rod cells that degenerate first when the retina detaches, followed by cone cells [25, 26]. Also, the effect of RD on cone cells is similar to that of rod cells.

Our model simulations and analysis suggest that when the retina detaches, formation of new discs completely ceases or is significantly reduced. Our results further suggest that some disposal mechanism for discs must exist even though the normal shedding and disposal process is completely disrupted due to the separation of the RPE from the NL. Since the ROS continues to degenerate during RD and eventually dies after some time, by setting a critical length *L*_*c*_ below which the death of rod cells occurs, our model predicts the survival time for a rod cell in a detached retina which gives the time frame within which retinal reattachment can restore vision. This time frame is found to be dependent on the disc removal mechanism. Finally, we identify a range of disc addition and removal rates that can result in cell death during RD.

## Mathematical model for ROS renewal

The first and only mathematical model developed to study the normal dynamics of ROS renewal, given by equations (1)-(3), is comprised of a system of ODEs that describes the time evolution of discs in various compartments of the ROS [24].

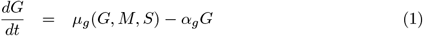

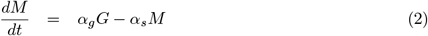

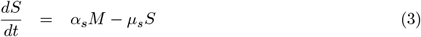

That is, the ROS discs are classified into three compartments: growth (G), mature (M), and shed (S). The growth compartment is located at the very base of the ROS. Discs in this compartment are still connected to the cell membrane, where they are open to the extracellular space and have not attained full disc diameter [27–29]. We assume that new discs are added to the G compartment at a disc addition rate *μ*_*g*_, which describes the average number of discs added to the ROS each day. The mature compartment (M) is located above the growth compartment and consists of fully developed discs detached from the OS cell membrane. We assume that discs from the G compartment mature into the M compartment at the rate *α*_*g*_.

This rate is governed by the rate of translocation of discs along the ROS. The discs are then shed from the mature compartment to the shed compartment at the shedding rate *α*_*s*_, being engulfed by the RPE. Finally, shed discs are disposed of by the RPE at disposal rate *μ*_*s*_.

Figure 1A shows the location of discs in each of the three compartments along the ROS, while Figure 1B is a compartmental diagram showing the transition of discs from one compartment to the other.

### Model assumptions

To explore the effects of RD on ROS dynamics, the original model proposed by the authors in [24] is expanded. In particular, to develop equations (1)-(3), the following assumptions for **normal ROS dynamics** were made:

1. The total length of the ROS constitutes only discs in the G and M compartments.
2. The number of discs in the ROS (i.e., *G+M* ) cannot exceed some upper bound *D*_*max*_. We assume this since the retina has a finite thickness such that ROS cannot grow without bound.
3. New discs are continuously added to the base of the ROS through membrane evagination at the disc addition rate *μ*_*g*_ [27–29].
4. The addition of new discs is faster when the ROS is short (during development), and slows down as the ROS approaches the maximum length (i.e., *G* + *M* → *D*_*max*_) [3]. As such, we define the rate of addition of new discs to be

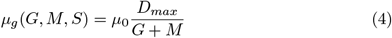

where *μ*_0_ is the minimum rate of addition of new discs.
5. Shed discs in the RPE are disposed of at the rate *μ*_*s*_. Although the exact disposal rate has not been quantified experimentally, it is a biological process that is known to occur [30]. Our mathematical model requires this disposal term so that the number of discs in the S compartment does not become too large. **To account for RD**, we make the following two assumptions:
6. Normal shedding, as defined to be the engulfment of discs at the distal end by the RPE [31], cannot exist during RD due to the separation of the RPE from the NL [20]. Since disc addition and shedding are the only two processes known to control the ROS length [32], the observed degeneration/shortening of the ROS during RD suggests the existence of some removal and disposal mechanism of ROS discs. To account for this removal process during RD, we multiply the shedding rate *α*_*s*_ by a dimensionless parameter

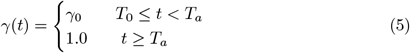

where *T*_0_ = 0 is the time of retinal detachment and *t* = *T*_*a*_ is the time of retinal reattachment as illustrated in Fig 2A. We assume that this parameter works to either increase, decrease, or keep fixed, the normal removal rate of discs. We assume this since there is no experimental data on such an alternative removal process. That is, we assume *γ*_0_ ∈ [0, 2] such that the alternative removal rate during RD is either less than (*γ*_0_ ∈ [0, 1)), equal to (*γ*_0_ = 1), or greater than (*γ*_0_ ∈ (1, 2]) the normal shedding rate.
7. RD prevents the supply of nutrients to the cells within the NL of the retina [33]. To capture the effect of RD on the addition of new discs at the base of the ROS, we assume that when the NL of the retina is detached from the RPE, the addition of new discs at the base is reduced by a certain factor due to the lack of supply of nutrients to the affected cells. In particular, we let

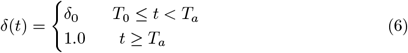

where *δ*_0_ ∈ [0, 1], as shown in Fig 2B, describes the severity of RD on the addition of new discs.

With the above-stated assumptions, and using the compartmental diagram shown in Figure 1B, the model we developed to help provide clarity to the precise effect of retinal detachment on the renewal of ROS is given by

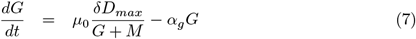

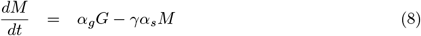

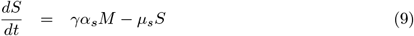

### Conversion from disk number to length

Experiments have been developed to measure ROS renewal dynamics in terms of ROS length [24]. Here we describe a method for converting disc number to length, which can help to compare our simulation results with experiments. As stated above in the model assumption, the total length of the ROS is proportional to the number of discs in the G and M compartments. As shown in Figure 1A, we let Δ_*T*_ be the average thickness of a disc and Δ_*s*_ the average disc-disc spacing. Then the approximate length of the ROS with *N* discs at anytime *t* (ie *N* (*t*) = *G*(*t*) + *M* (*t*)) is given by

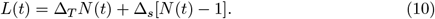

On the other hand, knowing the length *L*(*t*) of the ROS at any time *t*, we can estimate the number of discs using the relation

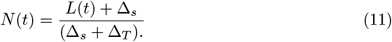

With the above relations, we perform simulations using the length associated with each of the three compartments. Going forward, we will denote the length associated with the number of discs in the G, M and S compartments by 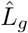, 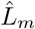 and 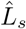 respectively. Therefore in terms of length, the model describing the renewal of ROS during RD is

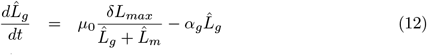

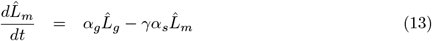

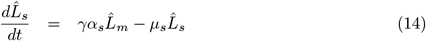

where *L*_*max*_ is the maximum length any ROS can attain and relates to the maximum disc number *D*_*max*_ by

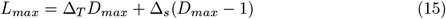

In simulating the model, we assume an initial condition of (*G*_0_, *M*_0_, *S*_0_) = (94, 627, 157) discs, which corresponds to initial lengths 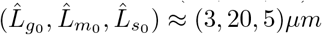 for mice. In addition, we assume that the initial time corresponds to the time of RD (if it occurs) and that the maximum length any mice ROS can attain is *L*_*max*_ = 30*μm* which in terms of disc numbers corresponds to *D*_*max*_ ≈ 940 discs. We choose this maximum value because the average length of mice ROS has been reported to be 23.8*μm* [34] and that of Royal College of Surgeons (RCS) rats to be 22.7 ± 2.4*μm* [35]. Table 1 lists the parameter values and references, as well as the units for the state variables and parameters.

**Table 1.** Parameter values. The table shows parameter values used in our simulation. The values for Δ_*T*_ and Δ_*s*_ correspond to those for mice [34]. The value for *μ*_0_ corresponds to RCS rat [35], the value for *α*_*g*_ was obtained from a study of zebrafish rods [24], while the remaining parameters *α*_*s*_ and *μ*_*s*_ were obtained through a parameter sweep where the model output (length of mature ROS) was compared to experimental data from mice [34].

| Quantity | Value | Unit |
| --- | --- | --- |
| $\hat{L}_g, \hat{L}_m, \& \hat{L}_s$ | - | $\mu m$ |
| $G, M, S$ | - | discs |
| $\Delta_T$ | $1.638 \times 10^{-2}$ | $\mu m$ [34] |
| $\Delta_s$ | $1.556 \times 10^{-2}$ | $\mu m$ [34] |
| $\mu_0$ | 2.2 | $\mu m/day$ [35] |
| $\alpha_g$ | 0.85 | $[1/day]$ [24] |
| $\alpha_s$ | 0.155 | $[1/day]$ estimated in this paper |
| $\mu_s$ | 0.3 | $[1/day]$ estimated in this paper |
| $\delta$ | $[0, 1]$ | No unit |
| $\gamma$ | $[0, 2]$ | No unit |

### Equilibrium and stability analysis

We start our model analysis by finding the equilibrium point(s) of the system. This will help us explore the long-term behavior/length of the ROS during retinal detachment. Since the ROS continuously degenerates during retinal detachment, we assume that there exists a critical length (*L*_*c*_) below which the rod cell dies. The equilibrium point(s) can help us determine whether or not the ROS length will eventually fall below this critical length. Before determining the equilibrium point(s) of the system and its stability, we first reduce the number of parameters in the model by nondimensionalizing the system. We scale the length associated with each of the three compartments 157 (equations (12) - (14)) by Lmax and time by 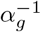 such that 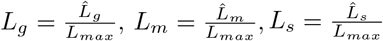, and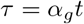, and arrive at

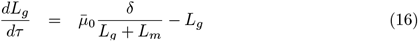

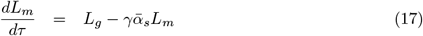

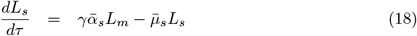

where 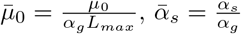 and 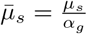. Setting the time derivatives in equations (16)-(18) equal to zero, we arrive at 2 equilibrium points 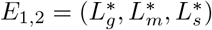 when *δ* ≠ 0, and a single trivial equilibrium *E*_0_ when *δ* = 0. That is, for *δ* ≠0, the equilibrium points are given by

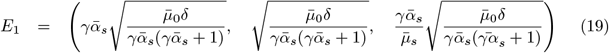

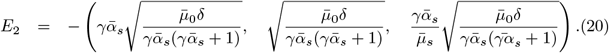

The equilibrium point *E*_2_ is not biologically feasible because each of the three state variables (*L*_*g*_, *L*_*m*_ and Ł_*s*_) corresponds to a non-negative length. When *δ* = 0, the two equilibrium points *E*_1_ and *E*_2_ coalesce into *E*_0_ such that

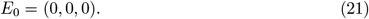

### Stability of the equilibrium point(s)

To determine the stability of the equilibrium points, we compute the Jacobian of the system defined by equations (16)-(18) which gives

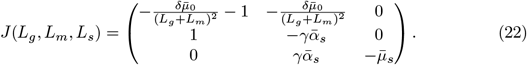

When *δ* ≠ 0 (i.e., making a substitution of *E*_1_ and then *E*_2_ into the Jacobian), the characteristic polynomial *P*_1,2_(*λ*) corresponding to the equilibrium points *E*_1_ and *E*_2_ is given by

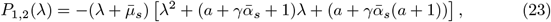

where 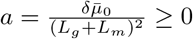. The equilibrium points *E*_1_ and *E*_2_ have the same eigenvalues given by

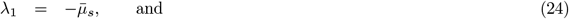

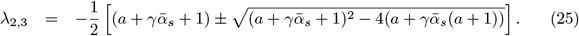

From equation (25), we observe that the real part of the square root term, 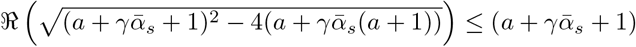. Thus, all the eigenvalues corresponding to equilibria *E*_1_ and *E*_2_ have negative real part which means they are both globally asymptotically stable. On the other hand, if RD completely stops the formation of new disks at the base of ROS such that *δ* = 0, the Jacobian of the system corresponding to *E*_0_ is

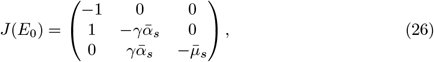

which has eigenvalues 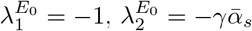 and 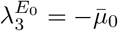. Thus, for *δ* = 0, if there is some disc removal rate such that *γ* ≠ 0, the equilibrium point *E*_0_ is also globally asymptotically stable. However, if *γ* = 0 such that there is no disc removal, the analytic solution to equations (16)-(18) is given by

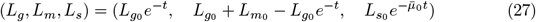

where 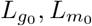 and 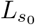 are the initial lengths corresponding to discs in the G, M and S compartments, respectively (i.e., at the time of RD).

## Result and discussion

Some studies suggest that RD halts the formation of new discs at the base of the ROS [19] but others suggest the contrary [16, 21]. Shedding, as understood by the engulfment of ROS discs by the RPE, ceases during RD due to the separation of the RPE from the NL [19, 20]. The precise effect of RD on the renewal of ROS is unclear. The ROS is observed to degenerate when the retina detaches [19, 20], and photoreceptor cells in a detached retina die after a certain time [16, 17, 21–23]. Thus, the mechanism responsible for the degeneration of ROS during RD, and those that dictate the survival time of rod cells in a detached retina are unknown. Here, we present the results of our model simulation and analysis to help answer these questions and provide clarity to the competing theories regarding ROS renewal. Resolving the controversies surrounding the renewal process and understanding the precise effect of RD on the renewal of ROS can help improve the existing therapy for RD and/or help devise new treatments for RD. Investigation of the survival time of a rod cell in a detached retina provides a time frame within which RD surgery can restore vision.

### Normal dynamics of a rod outer segment

Setting *δ* = 1 and *γ* = 1 in equations (12)-(14), we capture the normal dynamics/renewal of a healthy ROS. Here, we simulate the dimensional version of the system, given by equations (12) to (14), to better compare our results with experiments/clinical findings. Experiments on mice, rats, and frogs show that it takes on average 9-14 days for the entire length of the ROS to be renewed [13, 26, 36]. To highlight the *normal dynamics* of the ROS using our model, we assume that there is just one disc in the G-compartment initially with no discs in both M and S compartments, representing a developing rod cell. Fig 3 shows the length dynamics of all three compartments of the ROS and the total length (in red). Here, we used the average thickness of a disc (Δ_*T*_ ) and the disc-disc spacing (Δ_*s*_) for mice [34]. The predicted time to equilibrium is approximately 14 days, which is consistent with experimental findings.

**Fig 3.**
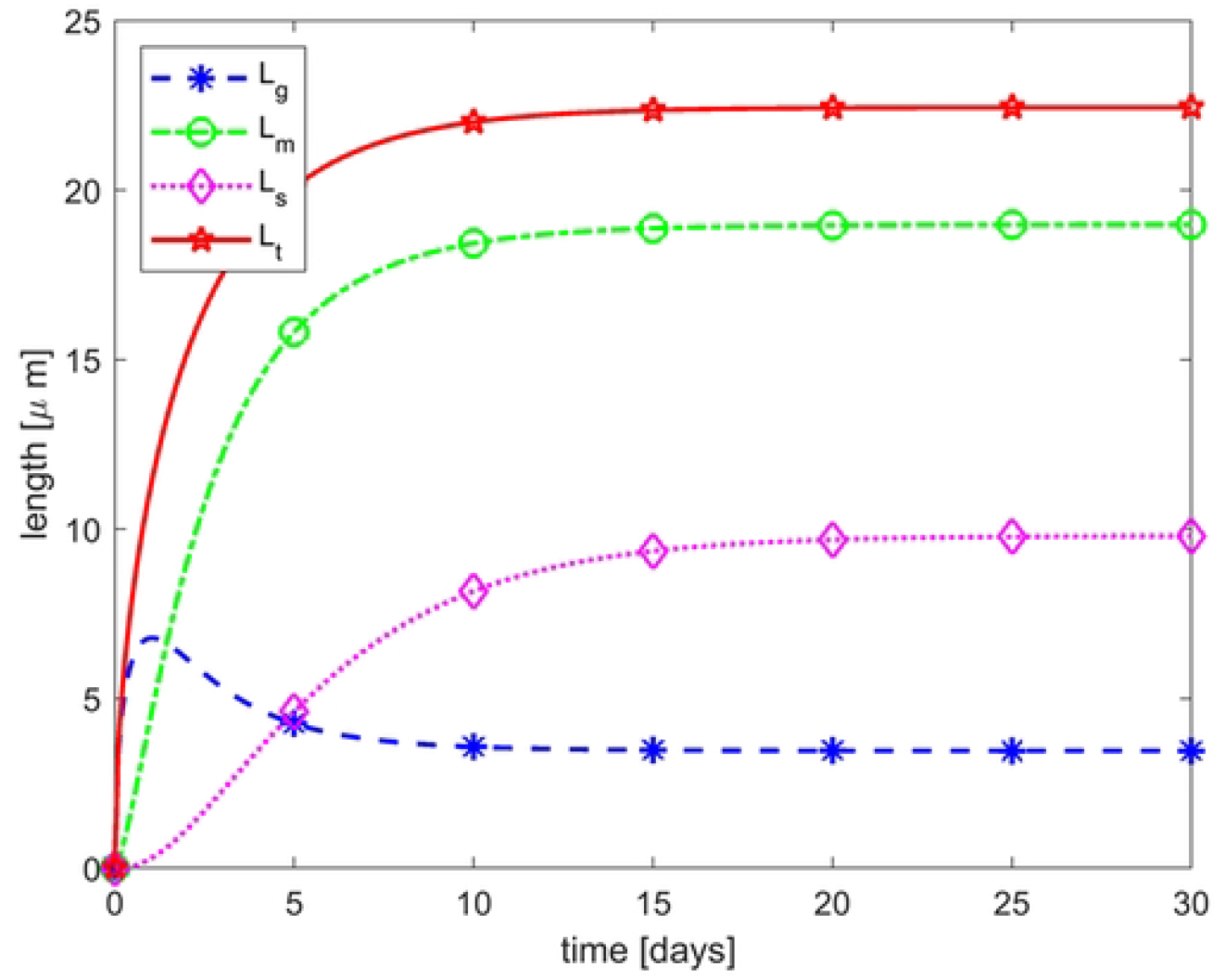
Normal ROS renewal. The figure shows the evolution of lengths associated with the G (blue curve), M (green curve), and S (magenta curve) compartments of a normal developing ROS, as well as the total ROS length (red curve where *L*_*t*_(*t*) = *L*_*g*_(*t*) + *L*_*m*_(*t*)). Initial condition is (*G*_0_, *M*_0_, *S*_0_) = (1, 0, 0) discs which translates to 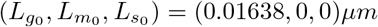 for mice. The time to equilibrium predicted by the model is approximately 14 days, consistent with experimental findings [13, 26, 36].

### Reduction in addition of new discs and complete suppression of disc removal (*δ <* 1 and *γ* = 0)

In general, it is observed that when the NL of the retina is detached from the RPE, the ROS degenerates/shortens [16, 19–21]. A degenerating ROS can be rescued to re-establish the equilibrium length if the NL of the retina is surgically reattached to the RPE before some **critical time** (*how long a rod cell can survive in a detached retina*) [16]. However, if the detachment lasts long enough, the rod cell might die and not be able to re-establish itself when the retina is surgically reattached to the RPE [17, 23, 37]. As stated previously, disc addition and shedding are the only two processes known to control the length of the ROS [32], and there is controversy surrounding whether or not disc addition persists or is halted. Although the normal shedding is prevented during RD due to the separation of the RPE from the NL, the observed degeneration of the ROS during RD suggests the possibility of some alternative removal mechanism. Here, we investigate whether or not disc addition can persist in the absence of an alternative removal mechanism. By setting *γ* = 0 to halt the disc removal process, we simulate the model for different reduced disc addition rates (i.e., *δ* ∈ (0, 1)), one of which is illustrated in Fig 4. Here, we assume that the normal addition rate is reduced by 98.64% (ie *δ* = 0.0136). Recall that the initial time of simulation corresponds to the time of RD. We observed that the ROS continuously increases in length throughout RD. We can infer from equilibrium point *E*_1_ in equation (19) that the long-term dynamics shown in Fig 4 is true for any *δ* ∈ (0, 1)). That is, since *L*_*t*_ = *L*_*g*_ + *L*_*m*_, when *δ* ≠ 0 lim_*γ*→0_ *L*_*t*_ → + ∞ indicating that once the alternative removal is suppressed, no matter how small the addition rate is, there will be an accumulation of discs which will increase the ROS length. This dynamics contradicts the experimentally observed degeneration/shortening of ROS during RD. Thus, our result suggests that disc removal cannot be halted if disc addition persists during RD.

**Fig 4.**
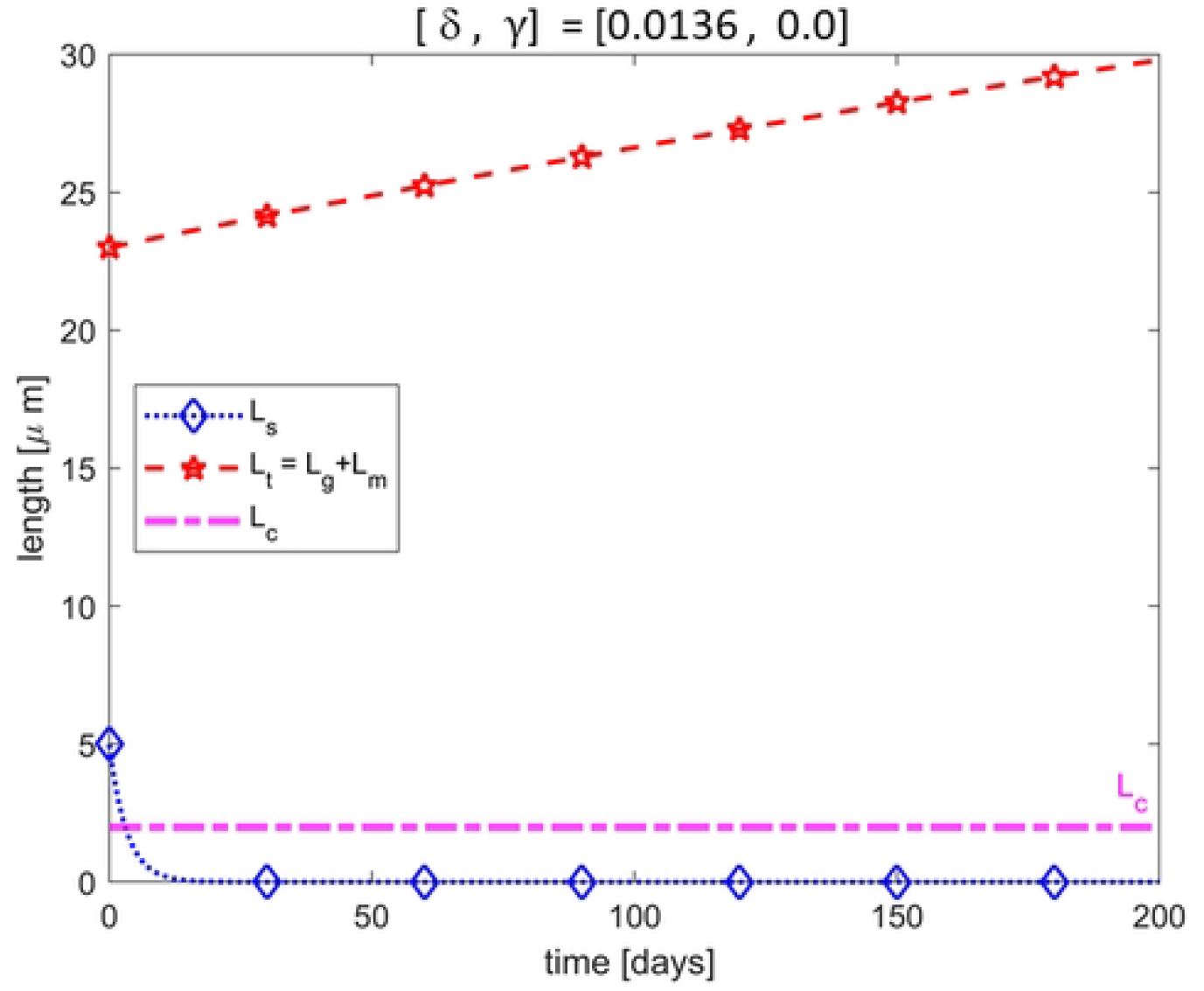
Addition without removal of discs. Illustration of the dynamics of a ROS during RD if the normal rate of addition of new discs is reduced, here by 98.64% (ie *δ* = 0.0136) while disc removal is halted completely. We observe an increase in the length of the ROS which contradicts the observed degeneration/shortening of ROS during RD.

### Halting of both addition of new discs and the removal of older discs (*δ* = 0 and *γ* = 0)

It is well established that the shedding of discs into the RPE ceases during RD. However, there could be some alternative removal mechanism taking place during RD. Here, we investigate whether or not the alternative removal mechanism as well as the formation of new discs can cease during RD. By setting *δ* = 0 and *γ* = 0, we assume that both the addition of new discs at the base and the removal of older discs at the top ceases when the NL of the retina is detached from the RPE. The analytic solution corresponding to this case is presented in equation (27). The phase portrait for this case is shown in Fig 5A in addition to the evolution of the total length of the ROS as well as the lengths corresponding to discs in S compartments in Fig 5B. These results show that the length of the ROS remains near-constant throughout RD. This result contradicts the experimentally observed degeneration of ROS that occurs during RD and inevitable cell death when RD is prolonged. Thus, the model suggests that the addition of new discs and the removal of older discs cannot both cease during RD.

**Fig 5.**
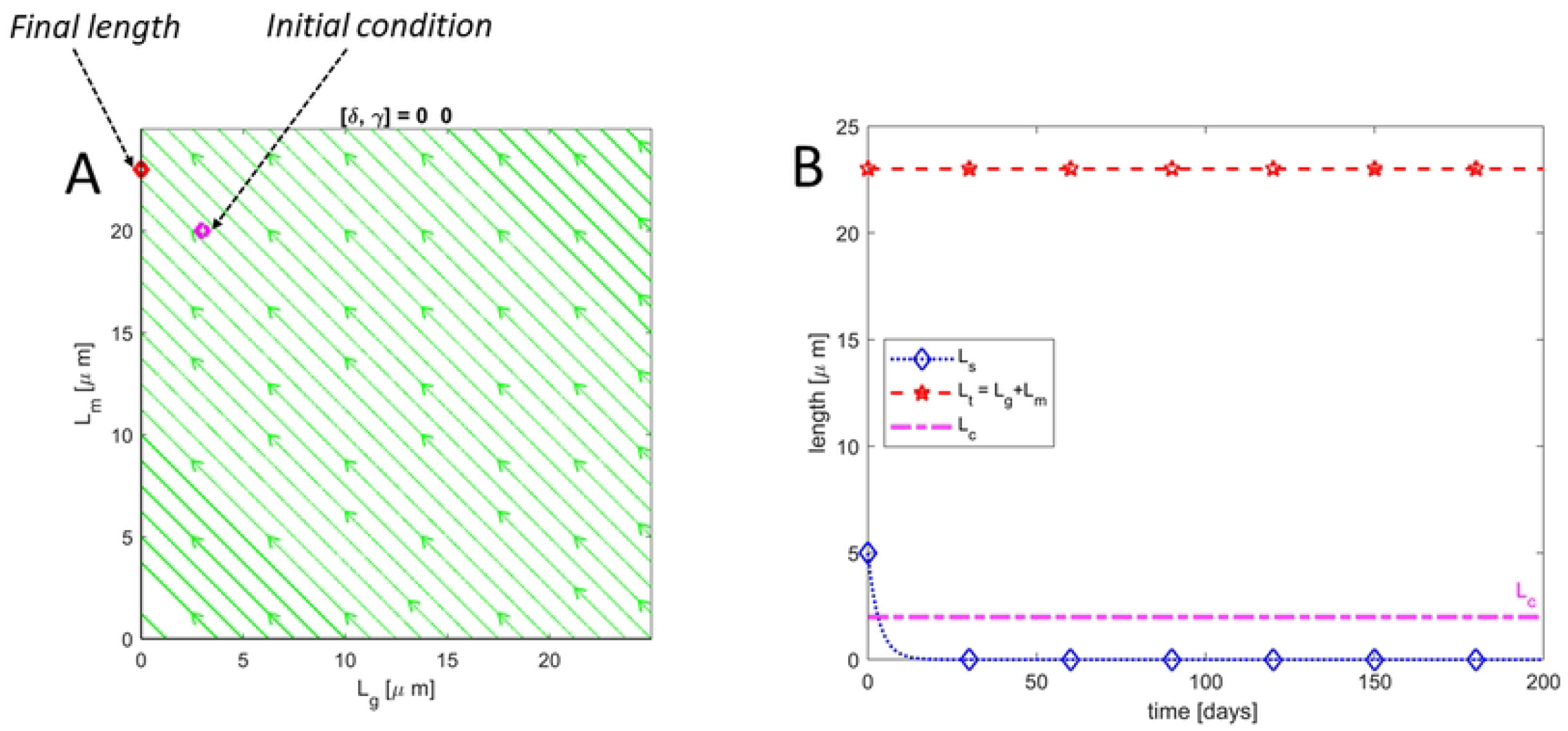
Neither addition nor removal of discs. **(A)** is a phase portrait in *L*_*g*_ − *L*_*m*_ plane and **(B)** shows trajectories for *L*_*t*_&*L*_*s*_ as well as the critical length *L*_*c*_ below which rod cell death occurs. Results show that the length of the ROS remains constant if both the addition of new discs and removal of older discs cease during RD thereby contradicting the observed degeneration/shortening

### Halting of addition of new discs and reduction in removal of older discs (*δ* = 0 and *γ* ≠ 0)

It is observed that RD leads to degeneration/shortening of the ROS and eventually cell death (ROS length falls below the critical length *L*_*c*_) if detachment lasts beyond a certain time *T*_*c*_. Here, we investigate whether or not RD prevents the formation of new discs but allows the removal of older discs. To do this, we set *δ* = 0 to prevent the addition of new discs at the base of the ROS and vary *γ* ∈ (0, 2) to allow for different removal rates. When *δ* = 0, the two equilibrium points *E*_1_ and *E*_2_ coalesce into *E*_0_ which is globally asymptotically stable. The output of our simulation, as shown in Fig 6, indicates that by halting disc addition and allowing the removal of older discs, the ROS continuously decreases in length towards zero. Thus, once the addition of new discs is halted, no matter how small the removal rate is, the ROS will eventually decrease below the threshold/critical length *L*_*c*_ thereby resulting in the death of rod cells. This result is consistent with the experimental finding by Kaplan, who suggests that RD inhibits the formation of discs or results in the abnormal assembly of new discs such that new discs are not able to configure themselves into the ROS [19]. Thus, our results suggest that RD completely stops the formation of new discs, a theory that is unclear in the literature as some studies suggest that disc addition persists during RD [16, 21]. In addition, since the RPE responsible for normal disc shedding is separated from the NL during RD thereby preventing the normal shedding process [19, 20], our model suggests that an alternate removal and disc disposal mechanism must exist during RD to explain the observed degeneration/shortening of the ROS and eventual cell death for prolonged RD.

**Fig 6.**
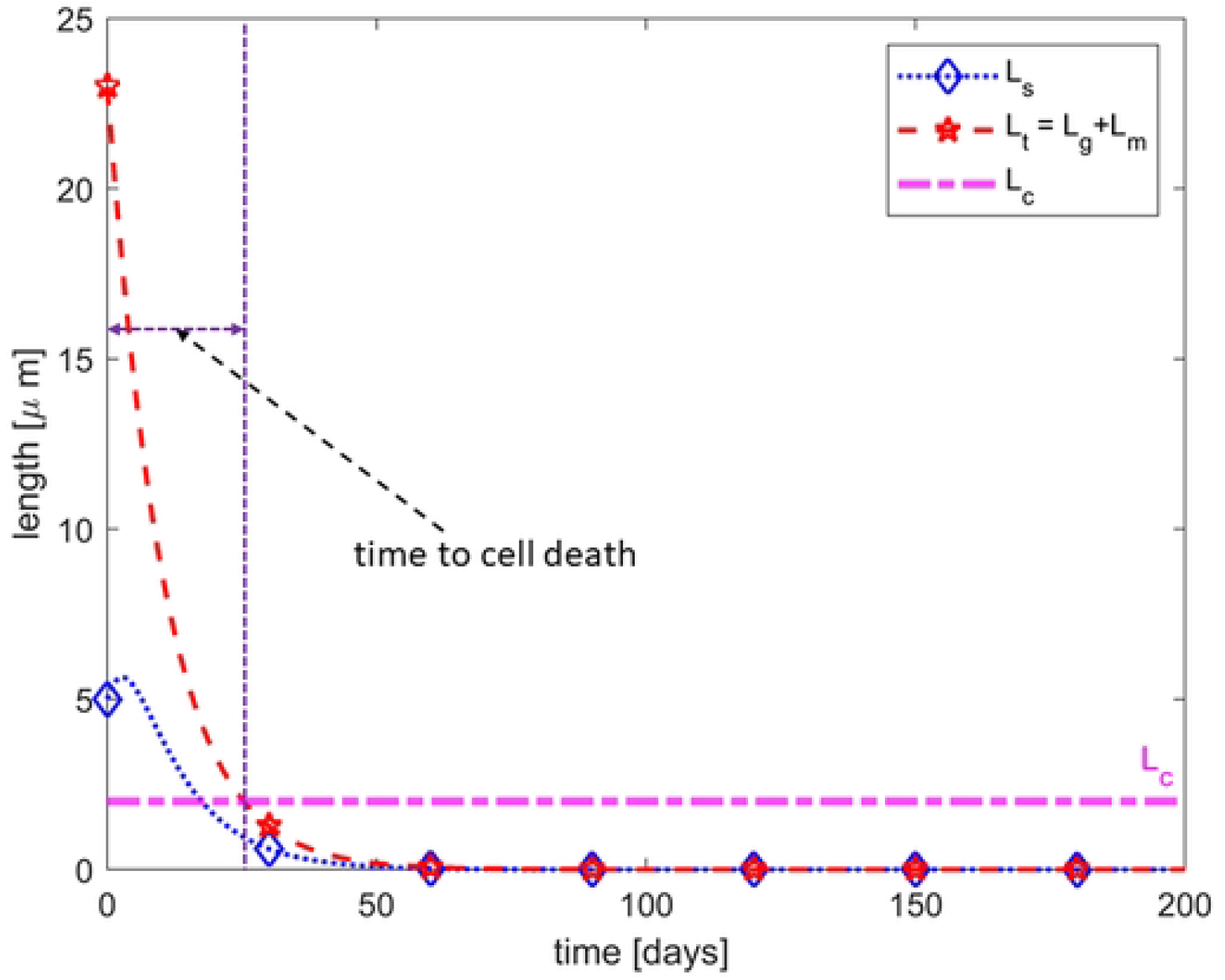
Removal without the addition of discs. Here the addition of new discs is halted (*δ* = 0) and there is a 37.5% reduction (*γ* = 0.625) in the normal removal rate. The ROS decreases below the critical length at approximately 25 days signifying cell death

### Reduction in both the addition of new discs and removal of older discs (0 *< δ <* 1 and 0 *< γ <* 1)

Here, we investigate the possibility of having both the addition of new discs as well as some removal mechanism at different rates during RD. Since RD prevents the supply of nutrients to the cells within the NL of the retina [33], we assume that the rate of addition of new discs cannot be more than the normal addition rate. Since no data exists on the alternative removal process, we propose that the alternative removal rate can be less than, equal to, or greater than the normal shedding rate. Varying *δ* ∈ (0, 1) and *γ* ∈ (0, 2), we respectively capture the different rates of addition of new discs at the base and removal of older discs at the top. As shown in Fig 7A, we note that during RD, if the rate of addition of new discs is significantly low (here 0.68% of the normal addition rate) the total length of the ROS decreases below the critical length *L*_*c*_ thereby resulting in cell death. However, if the addition rate is at least 1.3% of the normal addition rate, the total ROS length does not fall below the critical length (*i*.*e*., *rod cell does not die*) as shown in Fig 7B. This result supports the discovery by Kaplan [19], who suggested that RD either inhibits the formation of new discs or results in abnormal assembly of discs that are not able to configure themselves into the ROS. Since the formation of new discs occurs through membrane evagination at the ROS base thereby displacing already formed discs along the ROS [3, 28, 38], abnormal assembly of discs will result in very low disc addition rate leading to cell death when detachment last long enough. Thus, the model predicts that RD leads to a significantly low addition rate with some removal mechanism. The observed low addition rate in our model simulation supports the idea of an abnormal assembly of discs during RD.

**Fig 7.**
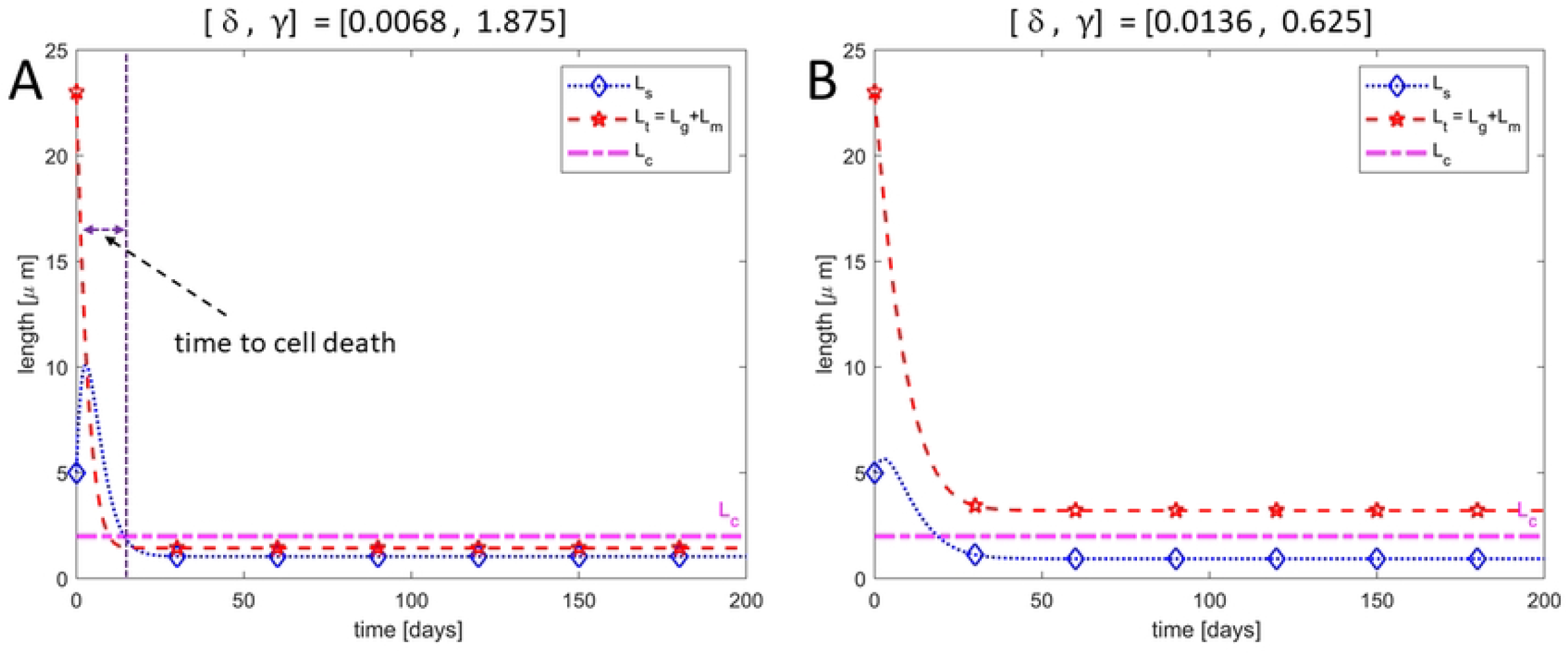
Formation and removal of disc. Illustration of the dynamics of a ROS during RD if both the addition of new discs and removal of older discs persist during RD (where disc addition is significantly reduced). In **(A)**, the normal rate of addition of new discs was reduced by 99.32% while the removal of older discs was increased by 87.5%. The ROS decreased below the critical length at approximately 10 days thereby resulting in cell death. In **(B)**, the normal addition rate was reduced by 98.36% while the normal shedding rate was reduced by 37.5% and the ROS did not decrease below the critical length (cell did not die).

We further determine the long-term behavior of the ROS during RD to help us predict when RD leads to eventual cell death (such that length falls below *L*_*s*_). Fig 8 A, B, and C are phase portraits in the *L*_*g*_ − *L*_*m*_ plane showing the stability of the equilibrium point *E*_1_ when *δ* ≠ 0 and Fig 8D describing the phase portrait when *δ* = 0. In all figures, we set *γ* ≠ 0 such that some removal mechanism exists since our model simulation and analysis suggest that an alternation removal mechanism is needed to explain the observed degeneration and cell death for prolonged RD. Here we see that when *δ* ≠ 0, decreasing *δ* (i.e., increasing level of severity of RD on the addition of new discs), the equilibrium point *E*_1_ moves closer to the origin where the total ROS length gets smaller but never reaches zero. Thus, if the formation of new discs persists during RD, the ROS cannot completely degenerate (i.e., ROS length cannot decrease to zero). However, depending on the rate of formation of new discs and the critical length *L*_*c*_ below which the cell can not survive, the ROS length may or may not fall below this critical length. When *δ* = 0, the two equilibrium points *E*_1_ and *E*_2_ coalesce to form a stable equilibrium point *E*_0_ at the origin. This means that no matter how small the critical length is, once formation of new discs ceases during RD, the ROS will always fall below the critical length signifying cell death if some removal mechanism of ROS disc exists.

**Fig 8.**
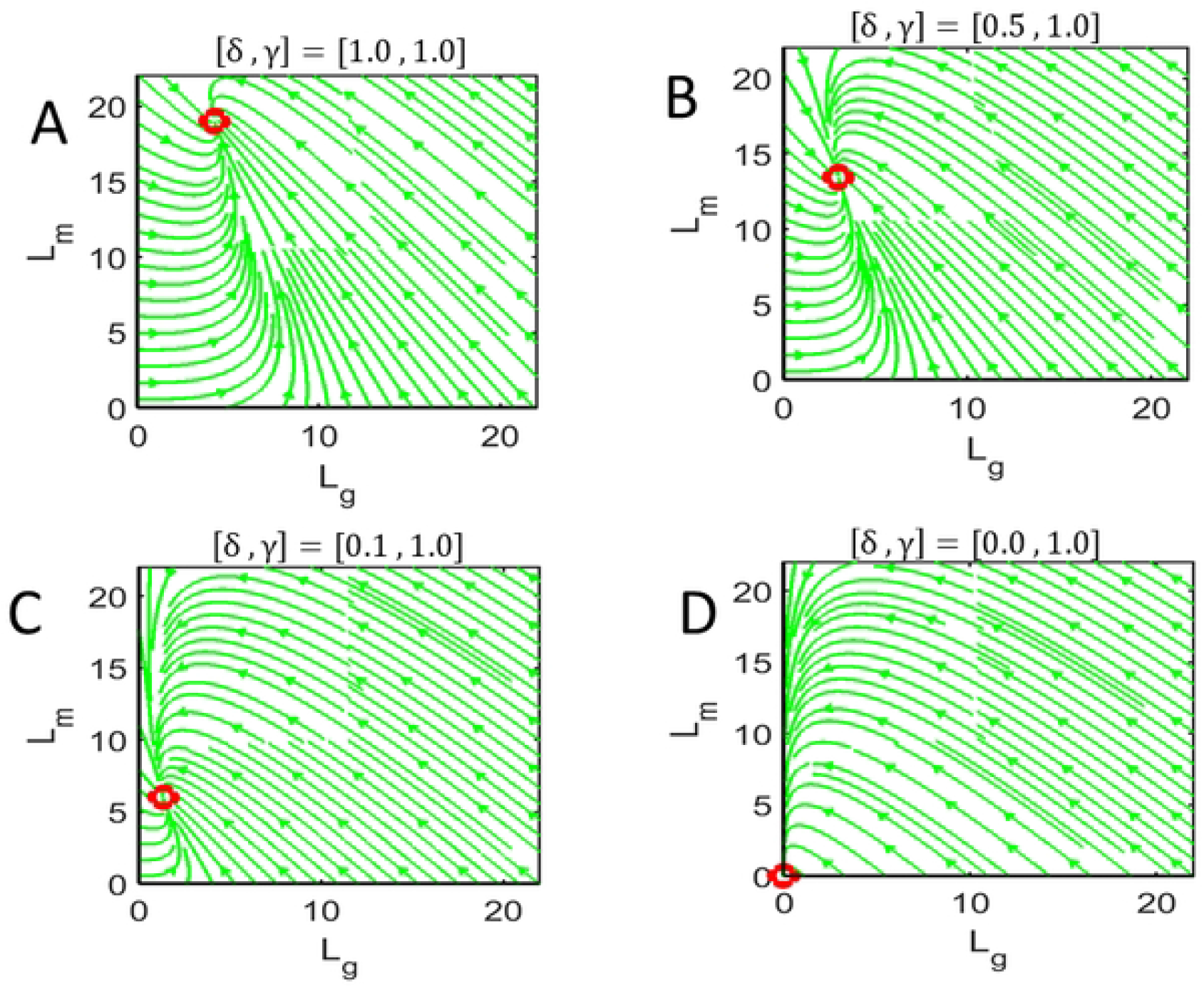
Phase plot. Phase portraits for varying disc addition rate *δμ*_0_ illustrating the effect of RD. Panel **A** shows a phase portrait for normal ROS dynamics (*δ* = 1 and *γ* = 1), panels **B** and **C** show an example of phase portraits during RD where we reduce the disc addition rate but keep the disc removal fixed (*δ <* 1 and *γ* = 1), and panel **D** illustrates a phase portrait when RD halts the addition of new discs (*δ* = 0).

### Criteria for unconditional ROS regeneration

Our model simulation and analysis results suggest that, for the ROS to shorten during RD, falling below some critical length *L*_*c*_ at critical time *T*_*c*_, **the rate of disk addition must be zero or close to it, and some disc removal mechanism must exist**. Here, we determine the range of values for the disc addition rate (*ν* = *δμ*_0_) and removal rate (*ε* = *γα*_*s*_) that can eventually drive the ROS below the critical length *L*_*C*_. As detailed in appendix S1 Appendix, the ROS length will fall below *L*_*C*_ if the addition and the removal rates satisfy the inequalities given by

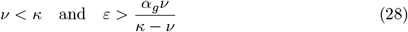

which we represent by the unshaded portion in Fig 9. This portion decreases or increases as we shift the vertical line *ν* = *κ* to the left or right respectively, where 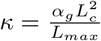. This result suggests that the size of the unshaded region is controlled by the critical length *L*_*c*_ and the disc addition rate *α*_*g*_ such that the smaller *L*_*c*_, the smaller the unshaded region, and vice versa. This means that the possibility of rod cell death is high if the critical length is large. The ROS will always regenerate upon reattachment of the RPE to the NL if the addition and removal rates satisfy the inequalities

**Fig 9.**
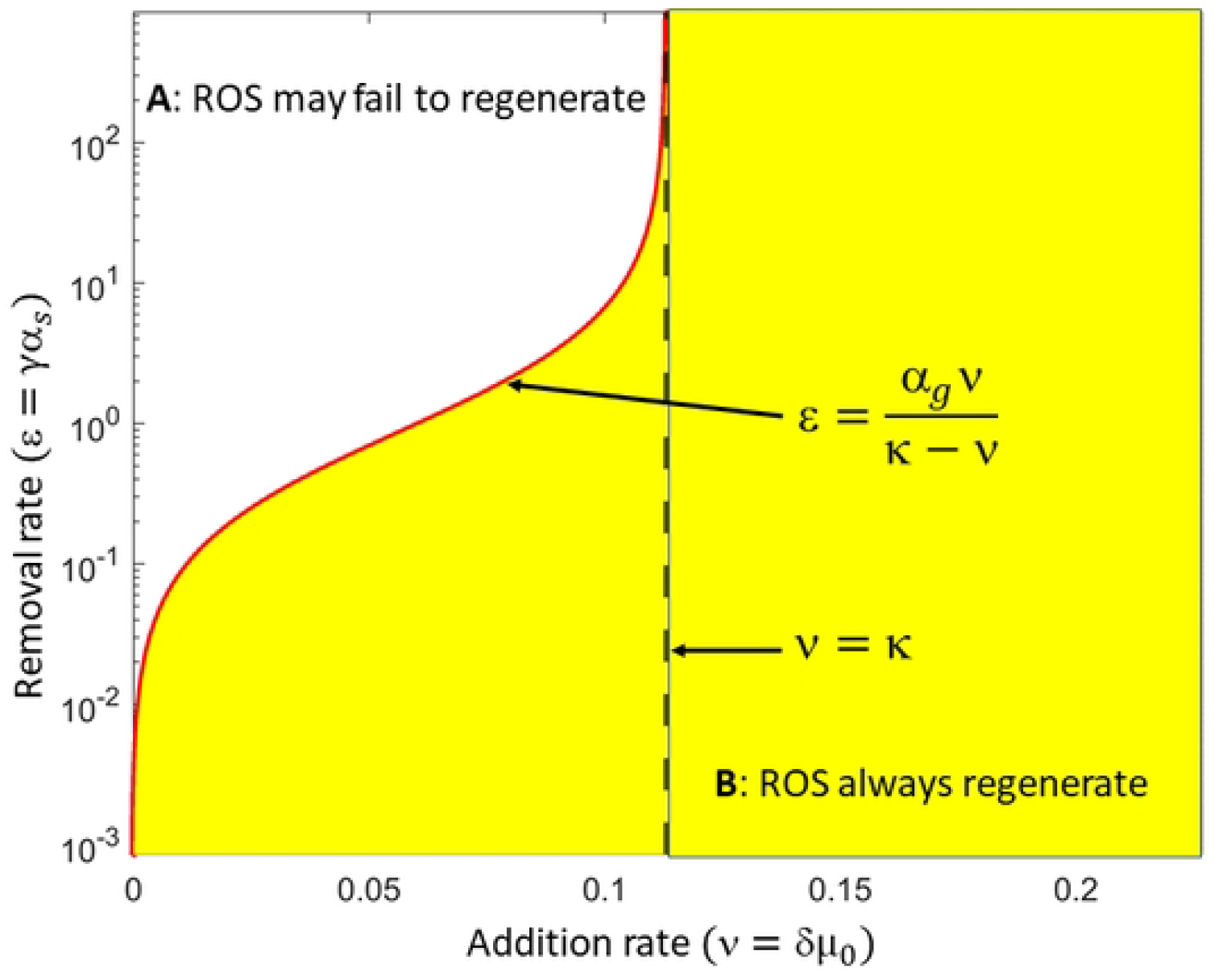
Domain for cell death. The unshaded portion labeled A in the figure shows the domain for the addition of new discs (*ν* = *δμ*_0_) and the removal of older discs (*ε* = *γα*_*s*_) during RD that will eventually drive the ROS length below the critical length leading to cell death. In the shaded region labeled B, the length of the ROS will always remain above the critical length *L*_*c*_, thus the rod cell cannot die and reattachment at any time will result in regeneration.

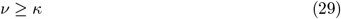

Or

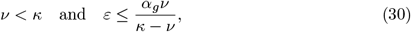

where the shaded portion in Fig 9 represents the range of values for the disc addition and removal rates for which the ROS remains above the critical length.

### Critical time to cell death

The recovery of sight after RD surgery depends on how long the retina has been detached [39, 40]. Here we investigate the time frame within which RD surgery can restore vision such that the length of the ROS is above the critical length *L*_*C*_. Biologically, this time frame corresponds to how long the photoreceptor (rod) cell can survive when the retina detaches. We investigate how the various parameters in the model influence the critical time *T*_*C*_ (the time at which the ROS falls below *L*_*C*_). We find that *T*_*C*_ depends on the disc removal rate such that the greater the removal rate, the smaller the critical time and vice versa. Thus if the removal rate during RD is higher than the normal shedding rate the critical time for reattachment is small. Fixing all other parameters (i.e., fixing *μ*_0_, *α*_*g*_, *α*_*s*_, *μ*_*s*_, and setting *δ* = 0) and varying the disc removal rate by varying *γ* ∈ [0, 2], Fig 10A shows how the critical times *T*_*C*_ varies. We observe that the *T*_*C*_ is more sensitive if *γ* ∈ [0, 0.5] but less sensitive if *γ* ∈ (0.5, 2]. In particular, there are large changes for the critical time for small changes in *γ* when *γ <* 0.5. For *γ >* 0.5, the critical time is small and changes only slightly as *γ* is increased. Similar analysis done for the maturation rate *α*_*g*_, as shown in Fig 10B, reveals that the critical time changes slightly when the disc maturation rate is small (*α*_*g*_ *<* 0.34) but has no effect on the critical time when the disc maturation rate is high (*α*_*g*_ ≥ 0.34). Figure Fig 10C, illustrates that the critical time is independent of variations in the disposal rate *μ*_*s*_, such that the critical time remains fixed. Based on these observations, the model suggests that the critical time at which the ROS can no longer regenerate is dependent and highly sensitive to changes in disc removal such that when *γ* is small the critical time is high but varies significantly for small perturbations in *γ*. For *γ* high, the critical time is small and varies only slightly for changes in *γ*. As the removal mechanism for discs during RD is not well understood (i.e., we do not know whether the removal rate is greater, equal to, or smaller than the shedding rate in normal ROS), our modeling results illustrate the need for better biological understanding of the disc removal process during RD so that we might establish the time frame within which RD surgery can successfully restore vision.

**Fig 10.**
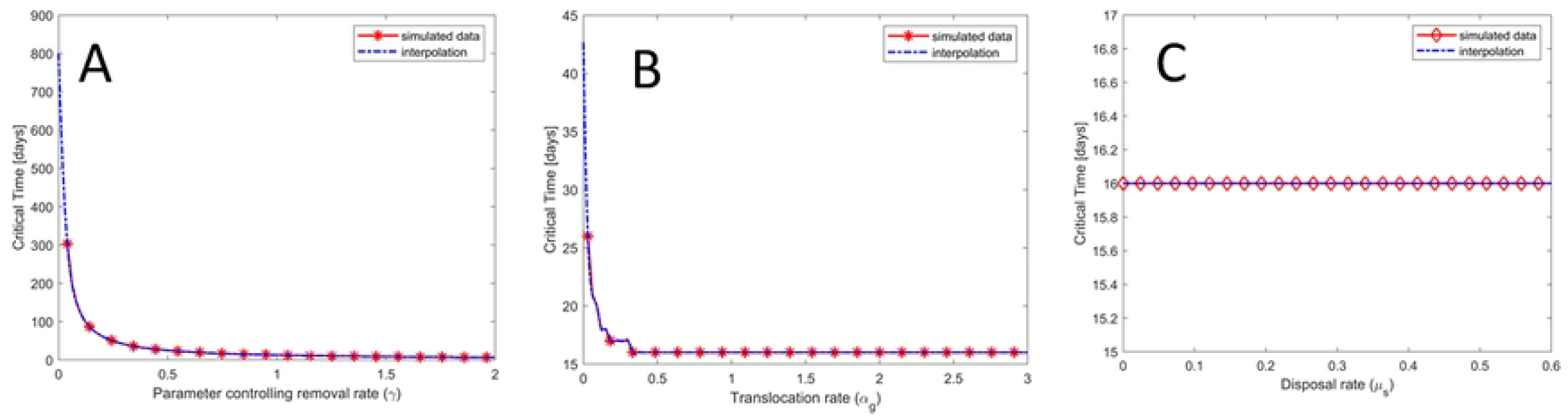
Sensitivity plot. The figure shows the dependence of the critical time on *γ, α*_*g*_ and *μ*_*s*_. In **A**, increasing *γ* (ie increasing the removal rate) decreases the critical time. In **B**, when *α*_*g*_ ∈ [0, 0.34) the critical time decreases with increasing *α*_*g*_ (*not as much as γ*) but does not affect the critical time when *γ* ≥ 0.34. In **C**, the disposal rate does not affect the critical time.

## Conclusion

We developed a mathematical model to describe the renewal process of ROS during RD. Using the output of our simulation, we were able to test competing hypotheses about the effect of RD on the addition of discs to the ROS. Our model suggests that disc addition is either halted or significantly reduced during RD [19]. Since we know that shedding, as defined in normal cells, can not occur as the RPE is removed from the NL [19, 20], our model also suggests that another removal and disposal mechanism must exist to get rid of discs at the distal end of the ROS to explain the observed ROS degeneration. Fig 11 describes our new idea (our modeling results) for how discs evolve in the ROS during RD (the old model is shown in red, the new is shown in black), where *ε* = *γα*_*s*_ represents the alternative disc removal rate during RD and *μ*_*r*_ describes a disposal rate. In addition to studying the effect of RD on the renewal process of the ROS, we also examined the time (the critical time *T*_*C*_) it takes for the ROS to decrease below a critical length *L*_*C*_ (which we assume results in cell death). As shown in Fig 10, we observed that the critical time is dependent on the rate at which ROS discs are removed, such that this time is very sensitive when *γ <* 0.5 but less sensitive when *γ* ≥ 0.5. Since we do not know if the removal rate is larger (*γ >* 1), equal to (*γ* = 1), or smaller than (*γ <* 1) the disc shedding rate in normal ROS. We also provided a quantitative description of how changes in the disc addition rate and the removal rate lead to eventual cell death during retinal detachment (the result is shown in Fig 10). This result indicates that the parameter ranges depend on the critical length (*L*_*c*_) below which cell death occurs and the maturation rate *α*_*g*_. This work sets the premise for a further experimental investigation into the threshold/critical length below which photoreceptor cells die, as well as the removal and disposal mechanisms of ROS discs during RD. Once these are established, the model can aid surgeons in understanding an approximate time frame or *window of opportunity* for which RD surgery will be successful.

**Fig 11.**
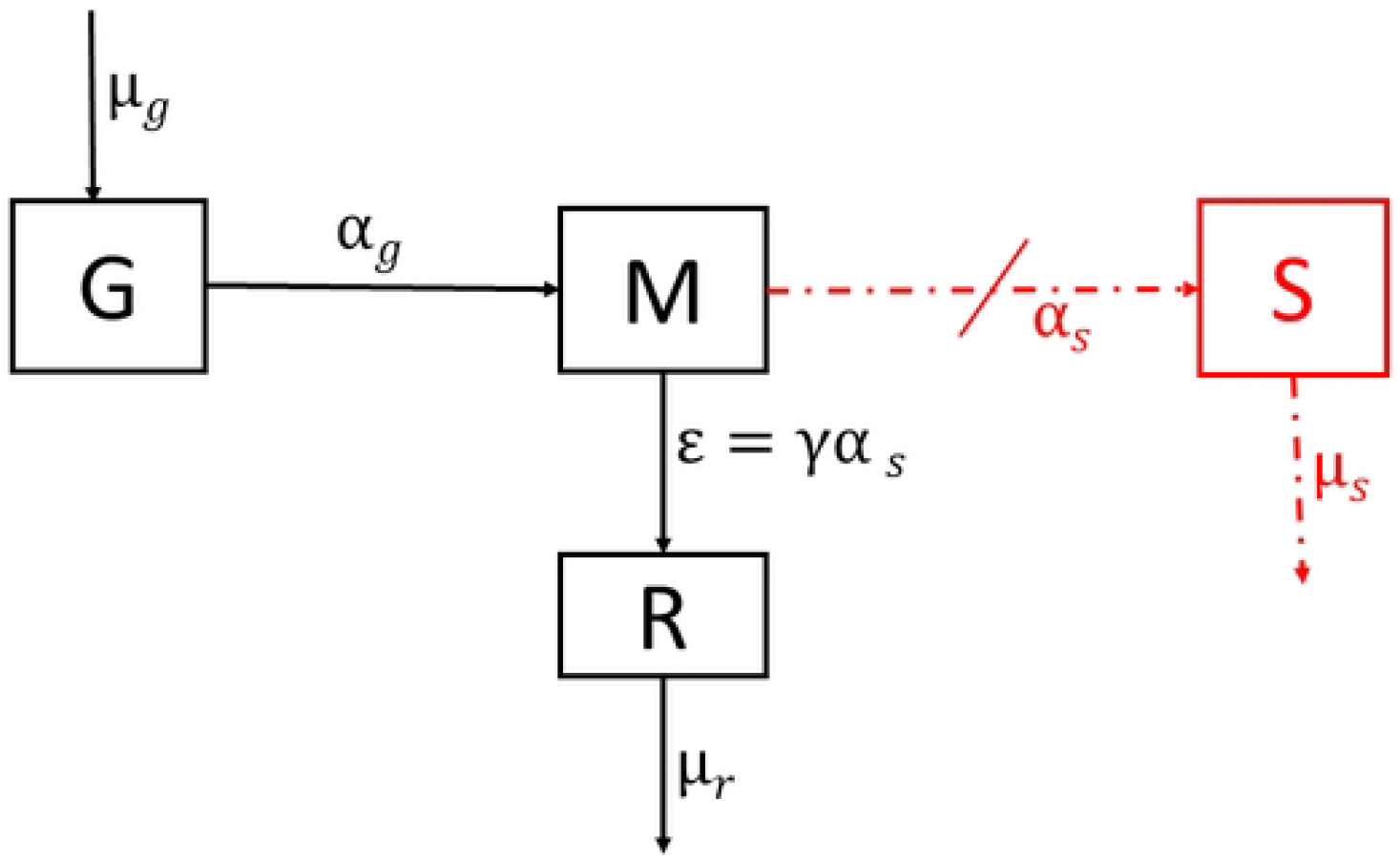
Proposed compartmental diagram. The figure illustrates a proposed mechanism that likely occurs during RD. The normal shedding as understood by the engulfment of discs at the distal end of the ROS by the RPE does not exist during RD due to the separation of the RPE from the NL. *ε* = *γα*_*s*_ is the proposed alternative removal rate of discs from the ROS during RD and *μ*_*r*_ the disposal rate.

The model proposed here is one of the first mathematical modeling frameworks to describe ROS dynamics during RD, where the authors have also used this modeling framework in [24] to describe the normal renewal process. As such, the modeling work completed here will be valuable in understanding other questions surrounding the ROS renewal process. In future work, we hope to look at extensions of this model to account for more complicated relations between the disc addition and shedding machinery as well as other factors regulating the renewal process. For example, we know that shedding is affected by the light-dark cycle [36, 41–44] and concentrations of certain chemicals such as dopamine and melatonin [45–47].

## Acknowledgments

This research was funded by the National Science Foundation under the Grant MCB-1951453. The authors would like to thank Dr. Abigail Jensen (UMass Amherst Biology Department) for engaging in helpful discussions that led to some important modeling considerations.

## Supporting information

**S1 Appendix.**

### Ranges for addition and removal rates that can lead to cell death

Retinal detachment leads to shortening of the ROS [48] and eventual cell death for prolonged detachment [39]. We assume that, for a rod cell in a detached retina whose outer segment (OS) is shortening, there exists a minimum length below which the rod cell is unable to survive, which we refer to as the critical length and denote by *L*_*c*_. Here, we determine the range of values for the removal rate (*γα*_*s*_) and the addition rate (*δμ*_0_) during RD that can drive the ROS below the critical length *L*_*c*_, leading to cell death.

For simplicity in notation, we denote the addition rate by *ν* = *δμ*_0_ and the removal rate by *ε* = *γα*_*s*_ and let 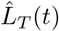 be the length of the ROS at any time *t*. Then, by adding the *L*_*g*_ and *L*_*m*_ components of the feasible equilibrium point *E*_1_ in equation (19), the equilibrium (total) length of the ROS is given by

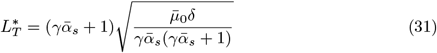

where 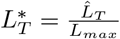 is a dimensionless quantity representing the total length of the ROS. For cell death to occur, we must have this length fall below the critical value such that

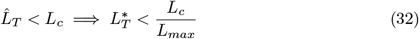

Substituting equation (31) into (32), squaring both sides and simplifying we arrive at

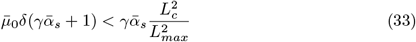

Recall from Section Equilibrium and stability analysis that 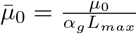 and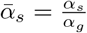. Substituting these into inequality (33) and simplifying gives

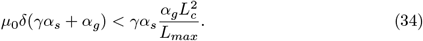

Denoting 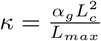 in inequality (34), we get

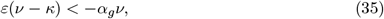

an inequality describing the relationship between disc addition and removal such that, if satisfied, will lead to rod cell death (i.e., total ROS length will eventually fall below the critical length *L*_*c*_). Since *ε, ν, κ, α*_*g*_ are all non-negative, inequality (35) holds only when

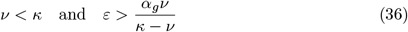

Fig 9 illustrates regions in *νε*-plane where the ROS will eventually fall below the critical length *L*_*c*_ signifying rod cell death or can not fall below *L*_*c*_ no matter how long the detachment lasts. In particular, if the rate of addition of new discs (*ν* = *δμ*_0_) and removal of older discs (*ε* = *γα*_*s*_) during RD satisfies the inequality in (36) (ie *ν* and *ε* falls within the unshaded portion of Fig 9), the ROS will eventually decrease below the critical length (*L*_*c*_) leading to cell death. On the other hand, if

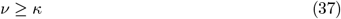

Or

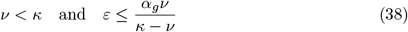

the ROS cannot decrease below the critical length *L*_*c*_. This means if the removal and addition rates of discs during RD fall within the shaded portion of Fig 9, the rod cell cannot die no matter how long RD lasts

